# Hierarchical transformations in sound envelope encoding differ across cortical layers

**DOI:** 10.1101/2025.09.09.674947

**Authors:** Chase A. Mackey, Yoshinao Kajikawa

**Affiliations:** Nathan Kline Institute for Psychiatric Research, Center for Biomedical Imaging and Neuromodulation, Orangeburg, NY 10962; New York University School of Medicine, Department of Psychiatry, New York, NY 10016

## Abstract

Amplitude-modulation (AM) is critical for the perception of complex sounds, and transformations in AM encoding may underlie primate-specific aspects of complex sound perception. The nonhuman primate model provides an opportunity to understand what circuit mechanisms generate hierarchical and interhemispheric transformations of AM encoding in different circuits across the cortical hierarchy. To address this, here we report the encoding of AM signals as a function of cortical layer and hemisphere in primary area A1 and the tertiary parabelt (PB) area of five NHPs. We presented AM noise and click trains to awake NHPs while recording from linear array multielectrodes spanning cortical layers. A1 typically encoded all AM frequencies (1.6-200 Hz) with high ( > 90%) classification accuracy, while the PB encoded lower (∼1.6-25 Hz) frequencies. The laminar gradient observed in A1 (Granular > Infragranular > Supragranular was inverted in PB (Supragranular > Infragranular > Granular), consistent with differences in thalamocortical input. Both areas displayed enhanced AM encoding in the left hemisphere, restricted to the supragranular layers. These results represent the first analysis of AM encoding in the PB, in which we identify circuits differing in their temporal sensitivity across the hierarchy, and suggest local supragranular neuronal populations contribute to hemispheric specialization.

## Introduction

Sound envelope fluctuations are a key sound feature that drives the perception of complex stimuli such as species-specific vocalizations and music (Ding et al., 2017; Henry et al., 2017; Rosen, 1992). Many previous studies have gained insight into how sound envelope encoding differs across the auditory pathway by recording neuronal responses to click trains and sinusoidally amplitude modulated (AM) tones and noise in nonhuman animals (Henry et al., 2016; Joris et al., 2004; Joris and Yin, 1992; Langner and Schreiner, 1988; Mackey et al., 2024a; Nelson and Carney, 2007; Rhode et al., 2010; Wang et al., 2008). These studies document a hierarchical progression across the pathway from a spike-timing based code, where spikes are aligned to individual cycles of the sound envelope (often measured using vector strength), to an average rate-code which is more prevalent at and above the inferior colliculus. By using measures of discrimination and classification performance derived from neuronal responses, studies of sound envelope encoding have explicitly addressed hypotheses about how information in different parts of the auditory pathway could contribute to sound envelope perception (Downer et al., 2017, 2021; Henry et al., 2016; Johnson et al., 2012; Mackey et al., 2024a; Nelson and Carney, 2007; Sayles et al., 2013; Yao and Sanes, 2021), with two of the critical findings being that 1) small populations of neurons in the brainstem, midbrain, and cortex encode information sufficient to account for detection and discrimination of amplitude modulation depth and frequency (Downer et al., 2021; Johnson et al., 2012; Mackey et al., 2024a; Nelson and Carney, 2007; Penikis and Sanes, 2023; Sayles et al., 2013; Shi et al., 2024), and 2) there are notable differences in encoding schemes in primary vs. secondary auditory cortex (Downer et al., 2017; Johnson et al., 2024). The use of small populations (empirical or simulated) has been a key aspect of these studies. In regions such as A1, single-neuron measures have suggested temporal information not to be present in sufficient quantities to account for the perception of AM based on, for instance, the population’s average vector strength alone (Johnson et al., 2012). However, subsequent studies utilizing decoding methods, which are agnostic to the presence of a rate vs. temporal code, have revealed that temporal information is preserved in the population (Downer et al., 2021). These findings informed the present study’s use of decoding methods on population-level activity measures.

In the study of AM encoding, the consideration of species is important for many reasons, and we focus on three here: 1) Primates (human and nonhuman) display greater perceptual sensitivity to AM than most animal models such as rodents and parakeets (Mackey et al., 2024a; Moody, 1994), 2) nonhuman primates (NHP) display more temporally precise AM encoding than rodents (Hoglen et al., 2018), and 3) primate auditory cortex exhibits a specific three-stage hierarchical anatomical organization, core-belt-parabelt, that provides the closest approximation of human auditory cortex available in animal models (Hackett et al., 2014, 2001). Previous studies of human auditory cortex report hierarchical progressions in speech encoding (Hamilton et al., 2021; Zion Golumbic et al., 2013) (e.g. Heschl’s gyrus vs. Superior Temporal Gyrus) and interhemispheric differences in the encoding of sound envelope (Boemio et al., 2005). Namely, the left hemisphere preferentially encodes higher frequency envelope fluctuations, while the right hemisphere preferentially encodes lower frequency envelope fluctuations; and Heschl’s gyrus encodes spectrotemporal speech features more precisely than the Superior Temporal Gyrus, which encodes higher level (e.g. semantic) information. The NHP model provides an opportunity to understand what circuits generate these transformations in encoding by characterizing AM encoding in different cortical circuits within area, and across the cortical hierarchy. For instance, it is possible that the hemispheric specialization reported previously is inherited from subcortical regions.

Laminar multielectrode studies (simultaneous recording across cortical layers) in auditory cortex have provided insight into the function of different intracortical and thalamocortical circuits in encoding temporal sound patterns involved in streaming (Banno et al., 2023; Fishman et al., 2004, 2001; Lakatos et al., 2013), intermodal audiovisual attention (Lakatos et al., 2009; Mehta et al., 2000; Mehta et al., 2000; O’Connell et al., 2015, 2014), and pitch perception of click trains (Steinschneider et al., 1998). Notably, previous studies in anesthetized cats have documented a laminar gradient in the precision of AM encoding (Granular > Supragranular > Infragranular), which provides a template to test in primary and higher order auditory cortex in primates (e.g. Atencio et al., 2009; Atencio and Schreiner, 2010). This is in part due to the fact that there is a dearth of NHP studies of AM encoding in the superior temporal gyrus, containing the higher-order parabelt region of auditory cortex; moreover, there are no published studies of AM encoding as a function of physiologically identified cortical layers in NHPs. Thus it is likely that studies of AM encoding with laminar resolution across the cortical hierarchy could yield insights into the thalamocortical and intracortical circuits utilized for AM encoding, and temporal sound pattern encoding more generally, in humans. To address this, here we report the encoding of AM signals as a function of cortical layer and hemisphere in the NHP primary and parabelt auditory cortex.

## Materials and Methods

All experimental procedures were approved by the Institutional Animal Care and Use Committee of the Nathan Kline Institute (NKI).

### Surgical procedures

Five nonhuman primates (Macaca mulatta, two female) C, H, Q, S, and W were implanted with headposts and recording chambers. Recordings from area A1 were done in chambers positioned to aim penetrations perpendicular to the supra temporal plane (Male W in the right hemisphere, female H in the left hemisphere, male S in the left hemisphere, Female C bilaterally). Three NHPs were also implanted with superior temporal gyrus chambers (female Q only on the left, female H only on the left, male W bilaterally). Surgical details for accessing primary and parabelt auditory cortex have been described in our previous publications (Kajikawa et al. 2017; Kajikawa et al. 2015). Specifically, based on presurgical MRI, the chambers were positioned to aim penetrations (during weekly acute recordings) perpendicular to the lower bank of the lateral sulcus, in the case of primary auditory cortex. For accessing the parabelt, presurgical MRI was used to target the superior temporal gyrus (STG) via multiple landmarks, which aided in positioning optimally over the STG: the superior temporal sulcus (∼20 mm above the ear canal and ∼10 mm above the zygomatic arch over AP 0-20 mm), the lateral sulcus, and the zygomatic arch (described extensively in Kajikawa et al. 2015).

### Identification of recording locations

Utilizing the surgical approach described above, we were able to access the areas of the lateral sulcus and superior temporal gyrus that corresponded to area A1 in primary auditory cortex, and the parabelt region of the superior temporal gyrus. We approached primary auditory cortex at angles nearly perpendicular to the layers. Recording sites were functionally defined as belonging to A1, as opposed to R, RT, or Belt, based on the sharpness of tuning, tonotopic progression, sensitivity to tones vs. noise, and granular layer onset latencies to BF tones (Lakatos et al., 2005a; Merzenich and Brugge, 1973; Rauschecker, 1997).

We have previously published on the identification of the parabelt, its potential tonotopic organization, and other aspects of its topography in the same NHPs reported in the present dataset (Kajikawa et al., 2015; Mackey et al., 2024b). As described in these studies, and described in *Surgical Procedures,* a combination of pre-surgical MRI and laterally angled recording chambers permitted us to penetrate perpendicular to the cortical layers during acute weekly recordings.

#### Stimulus parameters

While NHPs sat awake, AM noise or click trains were presented with randomized modulation frequency (AM noise) or presentation rate (clicks). For brevity, the term modulation frequency will be used throughout the manuscript to describe the presentation rate of the click trains. Modulation frequencies utilized in this study were 1.6, 3.1, 6.3, 8.8, 12.5, 17.5, 25, 36, 50, 71, 100, and 200 Hz. Signals were 640 ms long, 70 dB SPL, 100% modulated, and were ramped on/off using a linear ramp that was 5 ms in duration.

#### Electrophysiological signal processing

Data collection was conducted as described previously (Kajikawa et al., 2017; Mackey et al., 2024b). U-probes (Plexon) with a linear array of 23 channels (0.3– 0.5 MΩ at 1.0 kHz) at intervals of 100 or 200 micrometers between channels were used. Signals from all channels were amplified 10x, followed by 500x amplification before multiplexing and filtering. Each channel recorded field potentials (FP) (0.1–500 Hz) and MUA (500–5000 Hz using zero phase shift digital filtering, 48 dB/octave and rectified) simultaneously. When compared directly, the envelope MUA utilized here provides similar responses, and estimates of tuning, to thresholded multiunit spiking activity (Kayser, Petkov, Logothetis 2007; Super and Roelfsema, 2005; Shaheen and Liberman 2018), which supports the idea that our results are comparable to previous multiunit studies of click train and amplitude-modulation encoding. Moreover, a previous study found that AM encoding properties of single- and multi-unit clusters corresponded closely (Langner and Schreiner, 1988). Both envelope MUA and FP were sampled at 2 kHz and stored for offline analyses.

### Data Analysis

Current source density (CSD) was calculated as the second spatial, i.e. across-channel, derivative of the FP for each time point of the laminar CSD profile (Freeman and Nicholson, 1975; Kajikawa and Schroeder, 2011; Mitzdorf, 1985).

#### Phase-locking value

The phase locking value of CSD and MUA signals at individual channels were computed at each modulation frequency (1.6-200 Hz) using complex demodulation, equivalent to a Discrete Fourier Transform coefficient at a given modulation frequency (Lu et al., 2022; Tallon-Baudry et al., 1996). For each trial *k*, a signal (duration 𝑇), windowed by Hann function 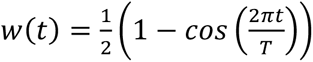, was convolved with a narrowband kernel, 𝑓(𝑡) = 𝑒^-2πift^ as 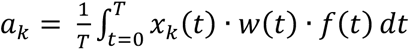. The resultant coefficient, 𝑎_k,_ is a complex, and its argument, φ_)_ = arg(𝑎_k_), depends on the timing of MUA fluctuation relative to the cycle of the band. The phase-locking value was derived from all *N* trials as 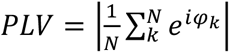. PLV is mathematically the same vector-length statistic used for vector strength of spike trains. In classical vector strength, the phases are taken at each spike time relative to the stimulus cycle. When computing PLV for continuous signals, the phase of the narrowband signal is calculated per trial by complex demodulation at the modulation frequency, yielding phases across trials. Both measures quantify the concentration of phase angles on the unit circle for each cycle of the stimulus from 0-1, with 1 being perfect synchrony, but PLV is better suited for our continuous CSD and MUA signals. And because we aggregate many cycles within each trial before taking the across-trial resultant, we avoid inflating sample size by counting cycles as independent; the degrees of freedom are the trials. This makes our PLV comparable to spike-based vector strength while remaining appropriate for continuous signals.

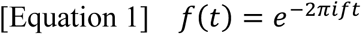

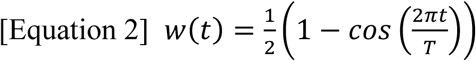

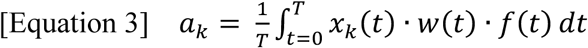

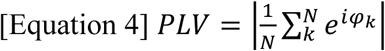

To statistically evaluate phase-locking, we constructed a random-phase null distribution (*n =* 1,000 permutations) that preserved the trial’s demodulated amplitude. We computed the observed amplitude-weighted vector, and estimated a *p-value* as the proportion of surrogates with a weighted vector greater than our observed weighted vector. We corrected for multiple comparisons across electrode channels using the Benjamini Hochberg method (Benjamini and Hochberg, 1995). For reference, we also computed corrected *p-*values from the more conventional Rayleigh’s test, which provided qualitatively similar significance across the dataset. To compute mean PLV across the dataset, only channels estimated to be within a layer-compartment, with significant PLV were utilized. The maximum frequency of significant phase locking was compared using a non parametric ANOVA.

#### Decoding Analysis and Mutual Information Estimation

To assess the decoding of stimulus identity from multichannel neural activity, we employed a multivariate classification and mutual information (MI) analysis pipeline based on linear discriminant analysis (“fitdiscr” in MATLAB; Statistics and Machine Learning Toolbox). This method fits a multivariate Gaussian model to each stimulus condition. The algorithm estimates the class-conditional means and a pooled covariance matrix, and constructs linear discriminant functions for each class. Each trial is then classified by assigning it to the class with the highest posterior probability, under the assumption of normally distributed features with equal covariance structure. This approach is equivalent to finding the linear projection that maximizes separation between class means while minimizing within-class variance. The analysis was conducted separately for each penetration site (i.e. recording session/experiment) and signal type (MUA or CSD) using custom-written MATLAB scripts. For each dataset, electrophysiological recordings were imported and epoched from -50 to 700 ms relative to stimulus onset and baseline corrected (-50 to 0 ms). Data used for decoding consisted of the time-domain signal from each trial and channel. For each layer (supragranular, granular, infragranular), only channels estimated to be within that cortical layer compartment were included for decoding. For each channel and decoding window, data were segmented into non-overlapping bins; unless otherwise specified, the decoding window was the entire baseline corrected 700 ms window after the stimulus onset, and we qualitatively inspected the pre-stimulus interval to ensure there was no difference in chance-level decoding; additionally, chance performance in the pre-stimulus interval is included in the figures. Linear discriminant analysis (LDA) classifiers were trained using fourfold cross-validation to predict stimulus condition labels from trial-wise data. A confusion matrix was computed for each fold, and the resulting matrices were averaged across folds to yield a single empirical confusion matrix per electrode channel, time window, and permutation. This calculation was repeated 1000 times to compute the 95% confidence intervals (CI) around “True MI” depicted in the results section. To compute chance performance, decoding analyses were repeated 1000 times with trial labels randomly shuffled. For each permutation and each decoding window, mutual information was computed from the resulting confusion matrix using the following formulation:

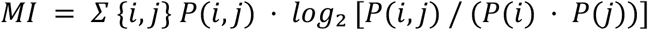

where P(i,j) is the joint probability of predicted and true labels, and P(i), P(j) are the marginal probabilities. MI was computed for both true and shuffled label conditions for all decoding time-windows, channels, and permutations. The resulting distributions were used to compute mean MI and 95% confidence intervals under both empirical and null conditions.

#### Statistical analysis

In all cases, criterion for significance was 0.05, and results requiring correction for multiple comparisons (e.g. the phase-locking values) were corrected via the Benjamini-Hochberg correction (see section on phase locking value). Linear mixed-effects model (LME) analysis was conducted using “fitlme” in MATLAB. We report the t-statistic given by fitlme, which is similar to the F-statistic often reported for multifactorial models. The t-statistic quantifies the strength and direction of the effect as the ratio of the coefficient estimate and its standard error. For mutual information analyses, only channels where the Mutual Information lower CI exceeded the shuffled (chance) Mutual Information upper CI were retained for statistical analysis. To account for the nested structure of the data (multiple channels per recording), we modeled MI as a function of three fixed effects: cortical layer (supragranular, granular, infragranular), area (A1 vs. parabelt), and hemisphere (left vs. right), including their two-way interactions. Individual channels were included as a random intercept term. Significant effects of layer, area, and hemisphere indicate a significant difference from the reference groups, which were the granular layer, area A1, and the left hemisphere. For best modulation frequency analysis, only channels where the bootstrapped lower CI exceeded the shuffled (chance) upper CI were retained for analysis.

In many nonhuman primate (NHP) neurophysiology experiments, including the present study, neural data are collected from many electrode contacts distributed across cortical layers and penetrations, across experimental sessions. In this context, the relevant statistical question is whether a given factor—such as cortical layer, area, or hemisphere—has a systematic effect on the measured neural variable (e.g., mutual information) across the population of recorded channels. This is distinct from testing whether different NHPs statistically differ from each other, which is a question that NHP studies are usually not adequately powered to address (Fries and Mars 2020). Accordingly, the degrees of freedom in our mixed-effects model reflect the number of auditory-responsive channels (with significant decoding accuracy) positioned in grey matter (confirmed using the CSD and MUA laminar response profile), recorded across sessions. For the CSD signal, we considered that the spatial resolution of the derived signal only permitted the isolation of a single sink (negativity), corresponding to a local synaptic current, in each layer compartment (supragranular, granular, infragranular). Thus we retained a single CSD channel per layer category for analysis, which corresponded to the largest sink (negativity) in that layer compartment, per recording. This limited our statistical power when analyzing CSD, and thus we mainly focus on multiunit activity when evaluating significance claims. For the multiunit activity signal, all auditory-responsive channels, with significant decoding accuracy, were retained for analysis, often totaling approximately 700-1000 channels across the dataset. For each analysis, the sample size can be inferred from the degrees of freedom, reported in its statistical table. For mixed effects models in MATLAB, the degrees of freedom correspond to the sample minus the number of fixed effects. Treating individual channels’ MUA as measuring different local populations is supported by our electrode spacing (100-200 micrometers), and the observation that neighboring channels displayed markedly different response characteristics (e.g. latency) and decoding accuracy.

## Results

Field potential and multiunit spiking activity were measured using linear array “laminar” multielectrode probes (see Methods) in response to amplitude-modulated noise, and click trains of different frequencies (i.e. repetition rates) in area A1 (AM noise, *n =* 42; Clicks, *n =* 63) and parabelt auditory cortex (AM noise, *n =* 88; Clicks, *n =* 84) across five rhesus macaques (*Macaca mulatta*); here, *n* refers to cortical sites, i.e. penetrations/experiments with the multielectrode probe. In all recordings, the electrode was angled to be nearly perpendicular to the cortical layers, and then spanned the layers during the recording (see Methods). From the field potential, current source density was calculated as the second derivative across electrode channels to mitigate volume conduction, and to estimate local transmembrane currents in specific cortical layer compartments with specific spatial resolution given by the interelectrode distance (100-200 micrometers). Using the frequently encountered activation pattern, characterized by an early onset, sharp L4 sink (negative CSD) co-localized with multiunit activity, followed by L2/3 and L5/6 CSD activation, individual electrode channels were assigned to the supragranular (L2/3), granular (L4), and infragranular (L5/6) compartments (Kajikawa et al., 2015; Lakatos et al., 2005b; Steinschneider et al., 1992). We have previously described how this layer-identification procedure can be used in the parabelt (see “The auditory spatiotemporal activation profile of the parabelt” in Mackey et al., 2024b; and see Kajikawa et al. 2015). We emphasize that application of this criterion to parabelt is subject to future anatomical validation due to different laminar patterns in primary sensory areas (e.g., thinner layer 4 in higher order areas such as the parabelt). However, the response properties observed provide some grounds for confidence in the layer assignment. Using this layer-identification procedure, we separated our data into local neuronal populations in the supragranular, granular, and infragranular layers for further analysis. This allowed us to utilize multiple channels of data per experiment, and to evaluate hypotheses about the function of neuronal populations in different cortical layer compartments. Figure 1 shows trial-averaged CSD and MUA responses to AM noise of different modulation frequencies. Figure 1A depicts the response to 3.1 Hz AM noise in a representative area A1 site, exhibiting an early (∼15 ms), sharp CSD sink (negativity, red in the color map) characteristic of the thalamorecipient granular layer. The multiunit activity derived from the same signal exhibits an early peak around in the same electrodes. The supragranular and infragranular CSD signals exhibited longer latencies, as expected, which resulted in relatively steady-state (as opposed to phase-locked) responses to the higher modulation frequency signals (Figure 1B-D). Figure 1E depicts an example parabelt site, where laminar response profiles displayed more diversity than area A1, which we have previously described (Kajikawa et al., 2015; Mackey et al., 2024b). A notable difference between the site in Figure 1G-H and Figure 1C-D is that the parabelt multiunit activity displays less phase-locking, though there is noticeable rhythmicity to the late MUA suppression that could contribute to discrimination between modulation frequencies (Figure 1G). Summary phase-locking values (see Methods) are depicted in Figure 1I-L. Across layers, similar phase-locking was observed, on average, in both the CSD and MUA signal. The phase-locking magnitude was consistent with previous reports using CSD and MUA in A1 (e.g. Steinschneider et al., 1998, 1993); and with single- and multi-unit measures of phase locking (e.g. Johnson et al., 2012; Yin et al., 2011); though the U-shaped trend in our analysis differed from previous studies. The parabelt differed noticeably from area A1 in that significant phase-locking was almost never observed above 10 Hz. We restricted calculation of the mean to frequencies at which more than 10 local neuronal populations (i.e. channels) displayed significant PLV (Figure 1I-L). The highest frequency at which significant phase-locking was observed was similar across layers, but differed between areas (Table 1). In area A1 (*n = 42* sites), MUA provided similar estimates of maximum frequency of PLV across layers . In the parabelt (*n* = 88), the maximum frequency of significant phase-locking was noticeably lower than A1 (Table 1). In A1, the differences between the layers in maximum frequency phase-locking were significant for the MUA signal, but not for the CSD signal ( Table 2). In the parabelt, the differences between the layers in maximum frequency phase-locking were significant for the CSD signal, but not for the MUA signal Table 2). The effect of brain area was significant after Benjamini-Hochberg correction for all layer categories for the CSD and MUA signals (Friedman’s test; χ^2^ (1) > 9.0, *p* < 0.01). While neuronal responses can be summarized using average amplitude or phase-locking measures, the underlying waveform is complex, particularly for continuous signals such as the CSD and envelope MUA depicted here. This motivated our use of decoding methods to assess information present in the response waveforms across our dataset.

**Figure 1.**
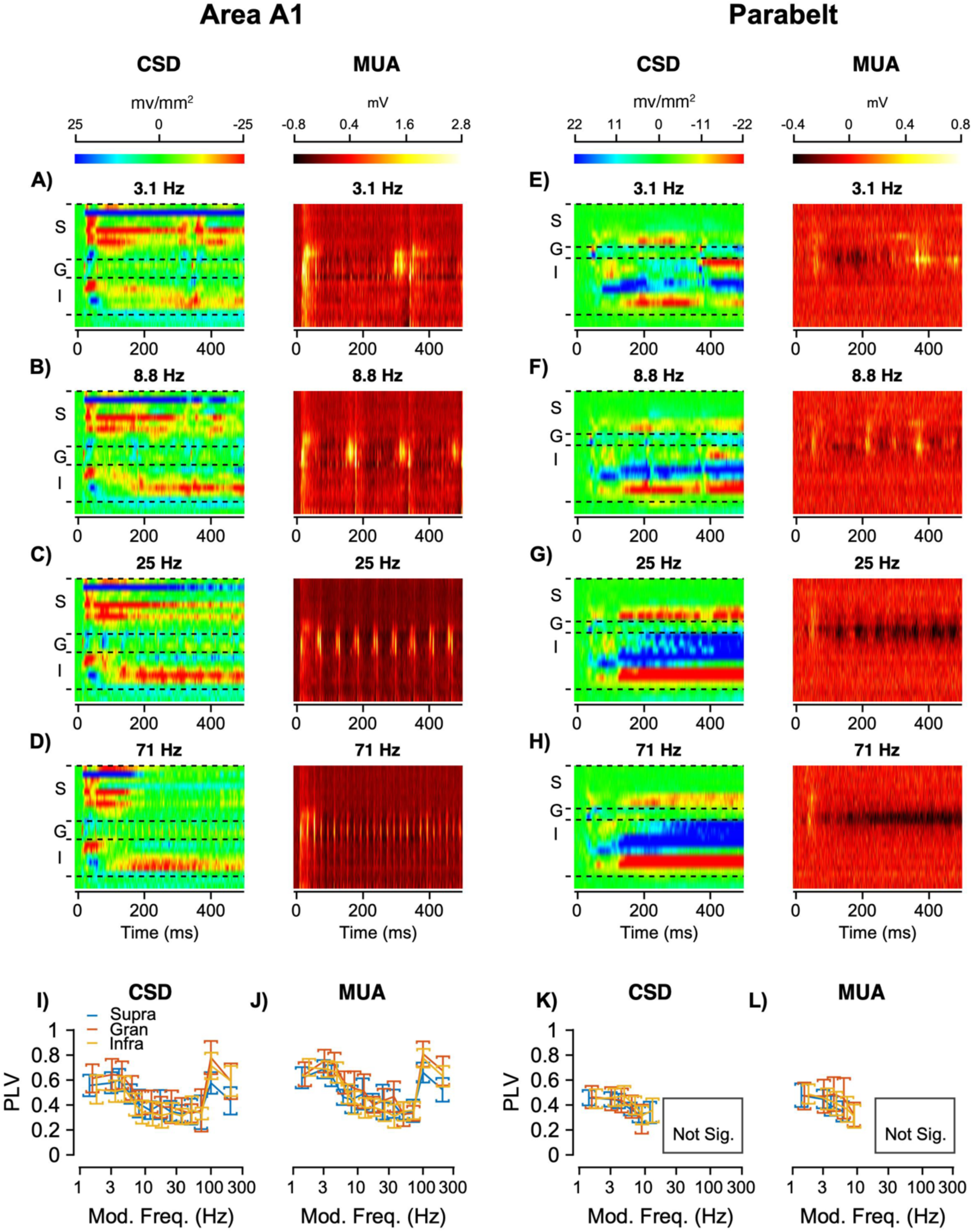
Representative laminar activity profiles in response to sinusoidally amplitude modulated noise of different frequencies, and phase-locking values across the data set. The left column (A-D) show trial-averaged responses (*n =* 50 trials) from a representative recording site in A1 to sinusoidally amplitude modulated (SAM) noise of different frequencies. The right column shows trial-averaged responses (*n =* 50 trials) from a representative site in the parabelt (E-H) to SAM of different modulation frequencies. These sites, and all of our recordings, were approached such that the electrode contacts were perpendicular to the cortical layers. A phase-locking value was computed for all recordings, which are displayed for each layer and signal type (CSD and MUA) for area A1 (I and J; mean and 95% CIs; *n* = 42 penetrations/sites) and the parabelt (K and L; mean and 95% CIs; *n* = 88 penetrations/sites). PLV was slightly jittered along the x-axis to enable visibility of the different data traces; in all cases, the modulation frequencies were the same (see Methods).

**Table 1.** Mean Maximum Phase-Locking Frequency by Area (A1, *n = 488* channels; Parabelt (*n = 258 channels*), Signal Type, and Layer.

| **A1** | **CSD** | Supragranular | 164.8 | 68.1 |
| --- | --- | --- | --- | --- |
|  |  | Granular | 172.5 | 59.8 |
|  |  | Infragranular | 161.6 | 70.3 |
|  | **MUA** | Supragranular | 152.7 | 69.9 |
|  |  | Granular | 169.1 | 66.2 |
|  |  | Infragranular | 170.3 | 68.4 |
| **Parabelt** | **CSD** | Supragranular | 50.5 | 54.0 |
|  |  | Granular | 42.2 | 56.1 |
|  |  | Infragranular | 50.3 | 54.9 |
|  | **MUA** | Supragranular | 35.5 | 64.9 |
|  |  | Granular | 22.9 | 38.5 |
|  |  | Infragranular | 26.9 | 36.9 |

Values represent mean (across sites) maximum frequency at which significant phase-locking was observed. CSD = current source density; MUA = multiunit activity; SD = standard deviation.

**Table 2.** Statistical Tests for Layer and Area Effects on Maximum Phase-Locking Frequency (A1, *n = 488* channels; Parabelt (*n = 258 channels*).

| A1 | CSD | Layer differences | $\chi^2(2) = 2.6$ | 0.26 (n.s.) |
| --- | --- | --- | --- | --- |
| A1 | MUA | Layer differences | $\chi^2(2) = 10.4$ | 0.006 * |
| Parabelt | CSD | Layer differences | $\chi^2(2) = 11.4$ | 0.003 * |
| Parabelt | MUA | Layer differences | $\chi^2(2) = 1.1$ | 0.59 (n.s.) |
| All layers | CSD & MUA | Area effect (Friedman's test) | $\chi^2(1) > 9.0$ | < 0.01 * |

All results are Benjamini-Hochberg corrected; n.s. = not significant; \* = significant ( $p < 0.05$ ).

### Classifying modulation frequency based on current source density and multiunit activity

To quantify how CSD and MUA encode information about AM stimuli, we utilized linear discriminant analysis (LDA; see Methods), and calculated Mutual Information (MI). MI is useful because it quantifies decoding accuracy across conditions, and the systematicity of decoding errors, as opposed to averaging accuracy across conditions. Figure 2 depicts the representative A1 and parabelt sites from Figure 1, with the LDA-derived confusion matrix for a channel in the granular layer for both sites, with MI values (right-hand panel) for all channels across layers (the y-axis). The color maps and MI plots were scaled the same across conditions and to emphasize the differences between area A1 and parabelt. Noticeably, area A1 exhibited nearly perfect decoding accuracy across conditions (Modulation frequencies). For example, the color map in Figure 2A indicates that most conditions were predicted correctly based on the granular channel CSD response in 100% of trials; and the granular channel MUA decoding was similar (Figure 2B). For each channel, MI was computed, which provides an information theoretic metric for stimulus information across different stimulus conditions (i.e. across the entire confusion matrix). 12 modulation frequencies were utilized, yielding a maximum possible MI value of about 3.5 bits (log_2_(12) = 3.5), which further emphasizes how accurate the site in Figure 2A-D is. Interestingly, though the overall decoding accuracy was similar, the laminar profile of MI differed between AM noise and clicks: supragranular MUA, for example, exhibited MI values not significantly different than the infragranular channels in response to clicks, while AM noise did not elicit such a strong response in the supragranular MUA.

**Figure 2.**
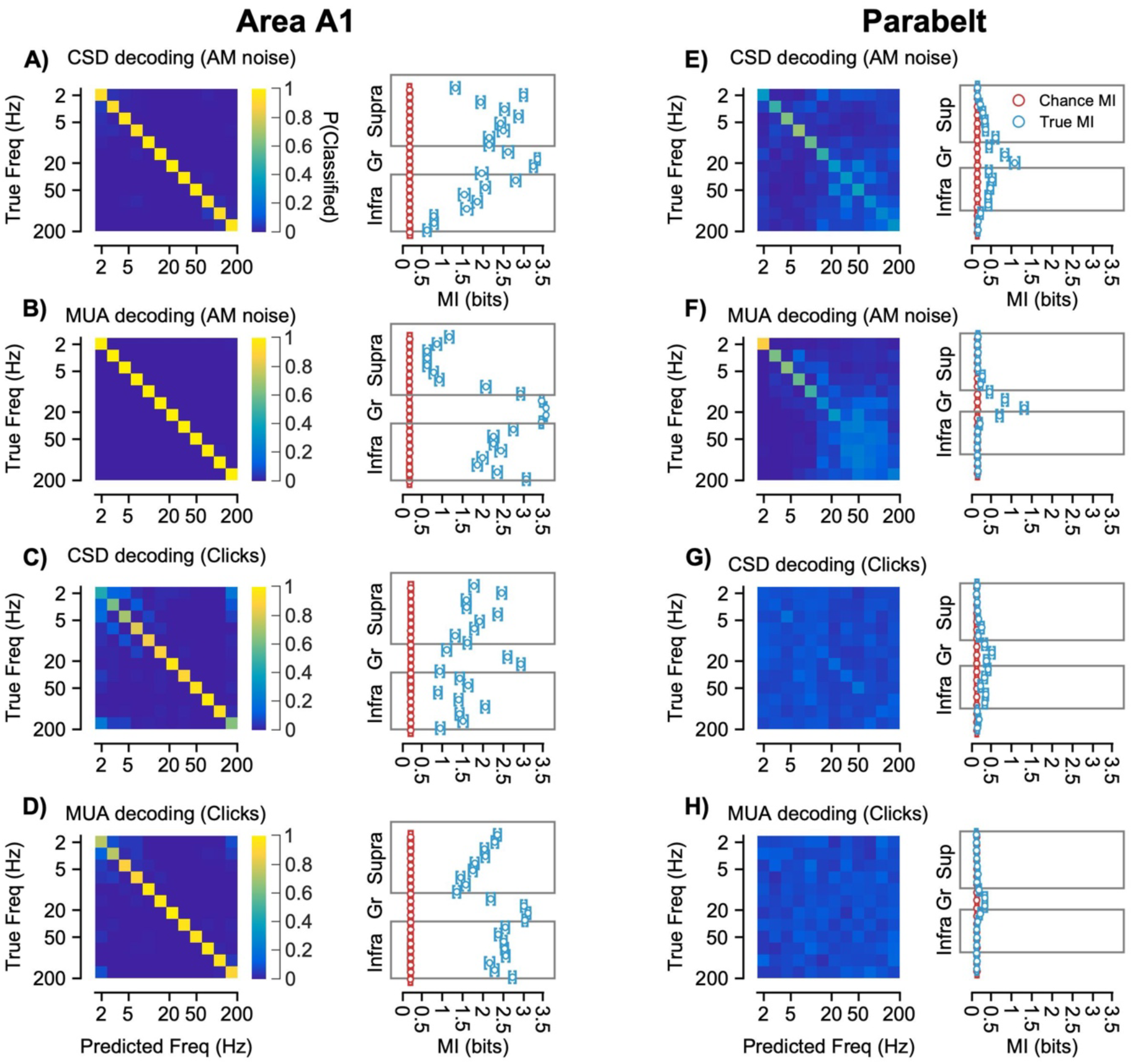
Representative example decoding results from AM noise and click-train evoked responses from the recording sites in Figure 1. The left panels depict the confusion matrices which quantify the decoding accuracy (0-1) in each condition for a channel in the granular layer. The predicted condition is indicated on the x-axis, and the true condition indicated on the y-axis. The color axis represents the proportion of classifications across 50 trials. The mutual information (MI) values for all channels are plotted on the x-axis of each panel on the right. Observed, or “true” MI was computed as described in the Methods, and Chance MI was calculated after shuffling the condition labels. For the example in area A1, the confusion matrices for a channel in the granular layer (layer 4) are shown in response to AM noise (**A** and **B**), and click trains (**C** and **D**), with corresponding mutual information values of all channels, which were calculated on the confusion matrices. Panels **E-H** depict the same analysis for the example in the parabelt.

Relative to area A1, the parabelt demonstrated reduced decoding accuracy and MI, likely due to the parabelt’s higher position in the processing hierarchy (Figure 2E-H). Though the granular channels exhibited the highest MI for this example, in the next figure we will show that the MI distributions were broader, i.e. there was more variability in the parabelt dataset than in area A1; thus, though the extragranular layers encoded very little information in this example, across the dataset they often displayed significant, above chance MI.

In the MI plots (right panels), channels are labeled as being in the supragranular (Supra), granular (Gr), or infragranular (Infra) layer compartments. Color maps are scaled 0-1, and mutual information plots are scaled 0-3.5 bits (the maximum possible MI with 12 stimulus conditions). Error bars represent 95% confidence intervals calculated across 1,000 permutations.

### Interhemispheric, interareal, and interlaminar differences in AM noise encoding

Across the dataset, we observed differences in AM encoding in different hemispheres, areas, and cortical layers. Figure 3 displays cumulative distributions of the MI values derived from CSD and MUA signals in the different hemispheres, layers, and cortical areas. Throughout the analyses presented here, the CSD and MUA will be shown, but statistical testing was reserved for the MUA results for reasons described in the Methods section.

**Figure 3.**
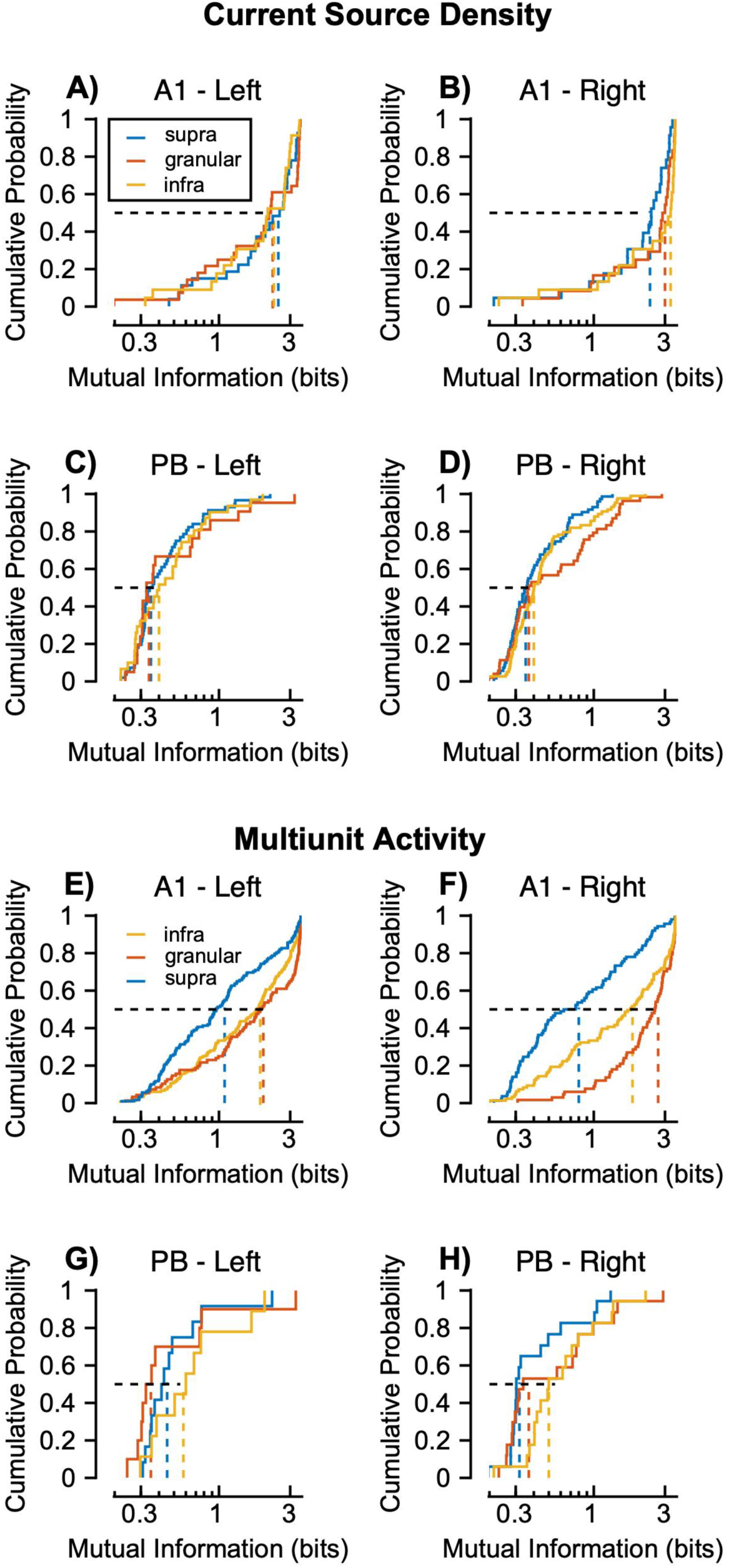
Cumulative distributions of mutual information values extracted from responses to AM noise across areas, layers, and hemispheres. Distributions of MI values derived from the CSD signal are depicted for left (*n =* 27, 28, 23 supragranular, granular, and infragranular channels, respectively) and right (*n =* 23, 24, 23 supragranular, granular, and infragranular channels, respectively) hemispheres of area A1 (**A-B**), and the left (*n =* 12, 10, 9 supragranular, granular, and infragranular channels, respectively) and right (*n =* 17, 20, 17 supragranular, granular, and infragranular channels, respectively) hemispheres of the parabelt (**C-D**). Distributions of MI values derived from the MUA signal are depicted for left (*n =* 131, 97, 154 supragranular, granular, and infragranular channels, respectively) and right (*n =* 86, 67, 99 supragranular, granular, and infragranular channels, respectively) hemispheres of area A1 (**E-F**), and the left (*n =* 21, 24, 25 supragranular, granular, and infragranular channels, respectively) and right (*n =* 21, 39, 38 supragranular, granular, and infragranular channels, respectively) hemispheres of the parabelt (**G-H**). Only above-chance values are displayed (see Methods). Dashed lines demarcate the median value of each distribution.

A mixed effects model was used to quantify the effects of each factor, and accounted for between-channel, and thus between-NHP variability (see Methods). MI was used as the dependent variable, while layer, area, and hemisphere were used as the independent variables (see Methods). Mixed-effects modeling revealed that mutual information (MI) was modulated by significant two-way interactions, which qualified the interpretation of main effects (Table 3). The reference groups for area, layer, and hemisphere were area A1, the granular layer, and the left hemisphere, respectively; i.e. all effects are quantified relative to these reference groups.

**Table 3.** Terms in a mixed effects model analysis of mutual information values extracted from MUA responses to AM noise (1.6-200 Hz) and click trains (1.6-200 Hz). The reference groups were A1, the granular layer, and the left hemisphere.

| Model: $MI \sim \text{Area} + \text{Layer} + \text{Hemisphere} + \text{Area} * \text{Layer} + \text{Layer} * \text{Hemisphere} + \text{Hemisphere} * \text{Area} + (1 \vert \text{Channel})$ | | | | |
| --- | --- | --- | --- | --- |
| Term: | AM noise ($df = 779, n = 776$ channels) | | Click train ($df = 997, n = 994$ channels) | |
|  | t-statistic | P-value | t-statistic | P-value |
| Right hemi (vs. L) | 2.2 | 0.025 | 2.96 | 0.003 |
| Supra. layer (vs. Gran.) | -5.2 | $3.0 \times 10^{-7}$ | -5.5 | $3.7 \times 10^{-8}$ |
| Infrag. layer (vs. Gran.) | -1.8 | 0.079 | -1.2 | 0.21 |
| Supra. layer*Right Hemi. | -3.2 | 0.001 | -3.45 | 0.0003 |
| Infra. layer*Right Hemi. | -1.6 | 0.09 | 0.2 | 0.81 |
| Supra. layer*Parabelt | 4.9 | $8.0 \times 10^{-7}$ | 2.7 | 0.0069 |
| Infra. layer*Parabelt | 3.3 | $8.0 \times 10^{-4}$ | 0.37 | 0.7 |
| Right Hemi.*Parabelt | -2.4 | 0.01 | -2.5 | 0.012 |
| Parabelt | -7.0 | $3.0 \times 10^{-12}$ | -11.8 | $2.0 \times 10^{-30}$ |

The first novel finding was that the hierarchical differences between area A1 and parabelt differed across layers. The main effect of area was reduced MI in the parabelt, though this differed across layers (Table 3). Specifically, decreased MI in the extragranular layers (relative to the granular layer) was observed in A1 (rightward shift in the orange traces in Figure 3 E-F; Table 3). However, this effect was attenuated in the parabelt (Fig. 3 G and H). This was quantified by a significant positive interaction of the extragranular layers with the parabelt, in the AM noise and click datasets (Table 3). The sign change in the *t-statistics* indicate that the laminar gradient observed in A1 (Granular > Infragranular > Supragranular) is inverted in the parabelt (Supragranular > Infragranular > Granular). We rotated the reference group from Granular to Supragranular, to Infragranular to confirm the inversion. Specifically, the granular reference group analysis is shown in Table 2 (*Supragranular*Parabelt*, *t = 4.9, p = 8*10^-7^; Infragranular*Parabelt, t = 3.3, p = 0.0008),* the Supragranular reference group analysis indicated the same effect (*Granular*Parabelt, t = -4.9, p = 8*10^-7^, Infragranular*Parabelt, t = - 2.27, p = 0.02),* and so did the Infragranular reference group analysis (*Supragranular*Parabelt, t = 2.27, p = 0.02, Granular*Parabelt, t = -3.4, p = 0.0008*). Thus, hierarchical transformations in AM encoding are not uniform across the auditory pathway, and may reflect differences in the information carried by thalamocortical and intracortical circuits. These findings will be considered in more depth in the Discussion section.

Previous work in human and nonhuman primate auditory cortex has indicated enhanced temporal processing in the left hemisphere (Boemio et al., 2005; O’Connell et al., 2015).. The present results suggest the supragranular layers play an important role in this asymmetry. The negative statistical interaction, *Supragranular Layers*Hemisphere*, confirms this quantiatively by indicating that the supragranular, but not infragranular layers, display left hemisphere enhancement (Figure 3 E-F; Figure 3 G-H; Table 3). The difference is indicated by the distributions of supragranular MI, which are visibly shifted to the right in area A1 in Figure 3E (supragranular median 1.1 bits) vs. Figure 3F (supragranular median 0.8 bits). Similar strength of the granular and infragraular interactions with the right hemisphere were observed for the MI values calculated based on click train responses (Table 3). These results highlight the importance of cortical layers when considering interhemispheric effects, and may aid in determining the circuit origin of hemispheric asymmetries, which will be further elaborated upon in the Discussion section.

A notable difference in the AM noise and click train conditions was that, in the infragranular right hemisphere parabelt dataset, less than twenty channels exhibited above-chance mutual information values. While this reinforces the finding that the left hemisphere and area A1 exhibit enhanced AM encoding, it suggests that click trains were not optimal for interareal comparisons. For this reason, the remaining analyses focus on the responses to AM noise.

### Topography of AM encoding in the parabelt

We next considered the hypothesis that caudal parabelt (CPB) exhibits enhanced AM encoding relative to rostral parabelt (RPB), based on CPB and RPB’s putative positions in the dorsal and ventral streams (Camalier et al., 2012; Hackett et al., 1998; Romanski et al., 1999). We quantified how mutual information about AM varied across the two-dimensional map of each NHP’s parabelt area using linear regression, which took the form “*Mutual Information ∼ X coordinate + Y coordinate*.” Mutual information did not significantly vary as a function of X (*t = 0.6, p = 0.5*) or Y coordinate (*t = -1.8, p = 0.07*). Moreover, neither X nor Y position significantly correlated with mutual information value (Spearman’s *R* < 0.2; *p > 0.05*). This can be seen in the representative example maps in Figure 4, which indicate that AM encoding did not systematically differ across the parabelt region in our dataset. Enhanced AM encoding would have greater temporal precision, i.e., preference for higher modulation frequency. We similarly tested whether best decoded modulation frequency displayed any topography in the parabelt, however, best modulation frequency did not significantly vary as a function of X (*t = 1.4, p = 0.17*) or Y coordinate (*t = 1.34, p = 0.21*). Moreover, neither X nor Y position significantly correlated with mutual information value (Spearman’s *R* < 0.4; *p > 0.05*). Finally, we similarly mapped the highest modulation frequency at which there was significant phase-locking, but found it did not significantly differ as a function of X (*t = 0.5, p = 0.62*) or Y coordinate (*t = - 0.837, p = 0.41*).

**Figure 4.**
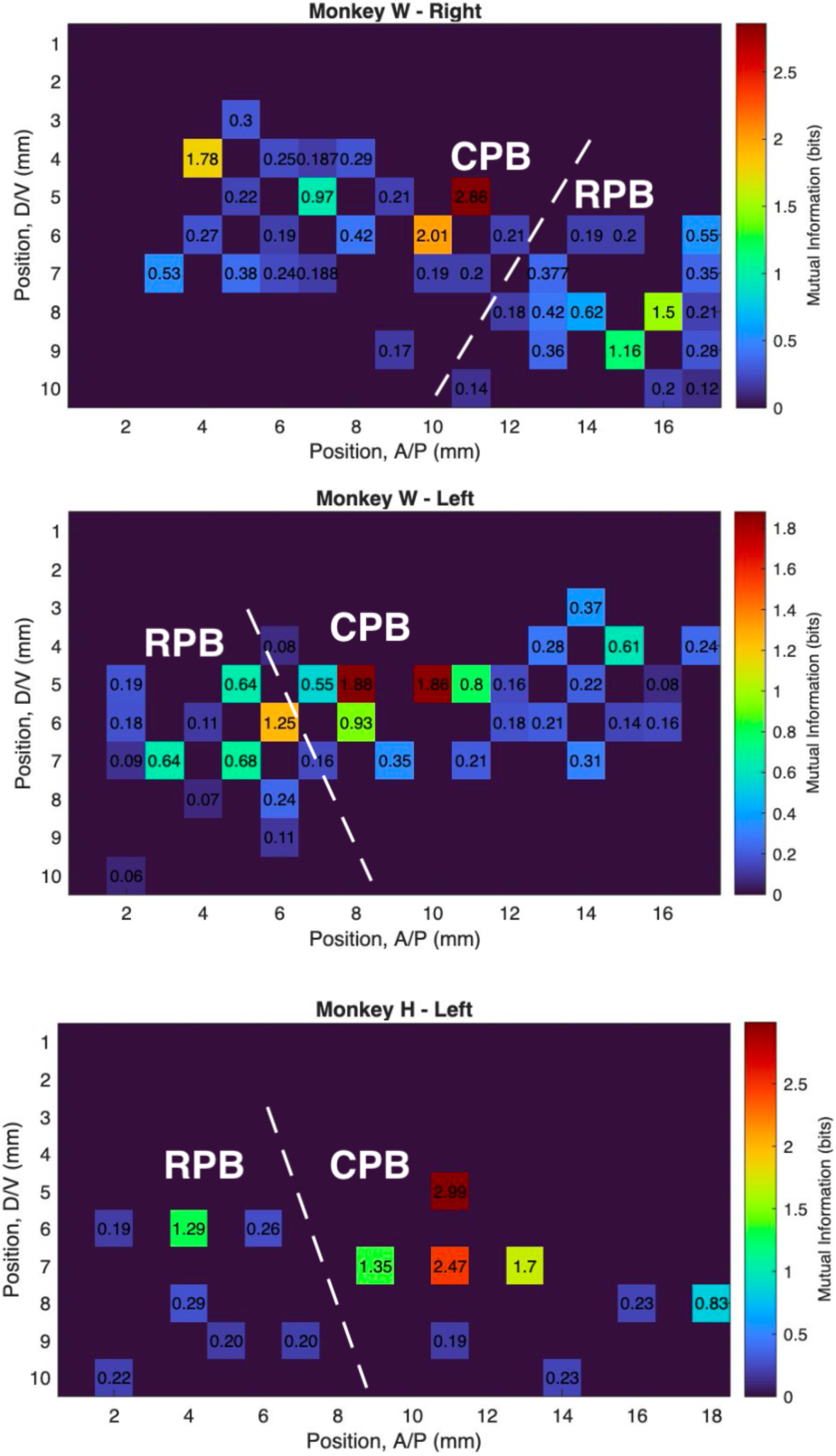
Topographic analysis of AM encoding in parabelt auditory cortex. Color maps depict mutual information values derived from MUA responses to amplitude-modulation, as a function of the two-dimensional maps across parabelt auditory cortex in three representative cases. As a result of the parabelt recording chamber placement over the superior temporal gyrus, the X and Y coordinates are arbitrary positions in a two dimensional grid with 1 mm resolution. The X coordinates refer to anterior-posterior (A/P) dimension, and the Y coordinates refer to the dorsal-ventral (D/V) dimension. The border between the rostral parabelt (RPB) and caudal parabelt (CPB) was determined using tonotopic progression.

### Preferred modulation frequency

Having characterized how mutual information differs across layers of A1 and PB, we next sought to understand how their preferred modulation frequency differed. Among MUA channels that significantly encoded at least one modulation frequency, we observed that the parabelt encoded lower modulation frequencies than A1 in a mixed effects model (*Best MF ∼ Hemisphere + Area + Layer (1|Channel);* Effect of Parabelt, *t = -4.4, df = 1536, p = 9*\*10^-6^). Best modulation frequency appeared to be higher in the infragranular layers of area A1 (Figure 5A), but this trend was not present in the parabelt dataset (Table 4; Interaction of Infragranular layers with Parabelt). Regarding hemispheric differences, the right hemisphere trended towards encoding lower modulation frequencies, and there was a significant positive interaction of right hemisphere with infragranular layers, which indicates that the infragranular layers encoded higher modulation frequencies than would be expected based on the right hemisphere main effect (Table 4).

**Figure 5.**
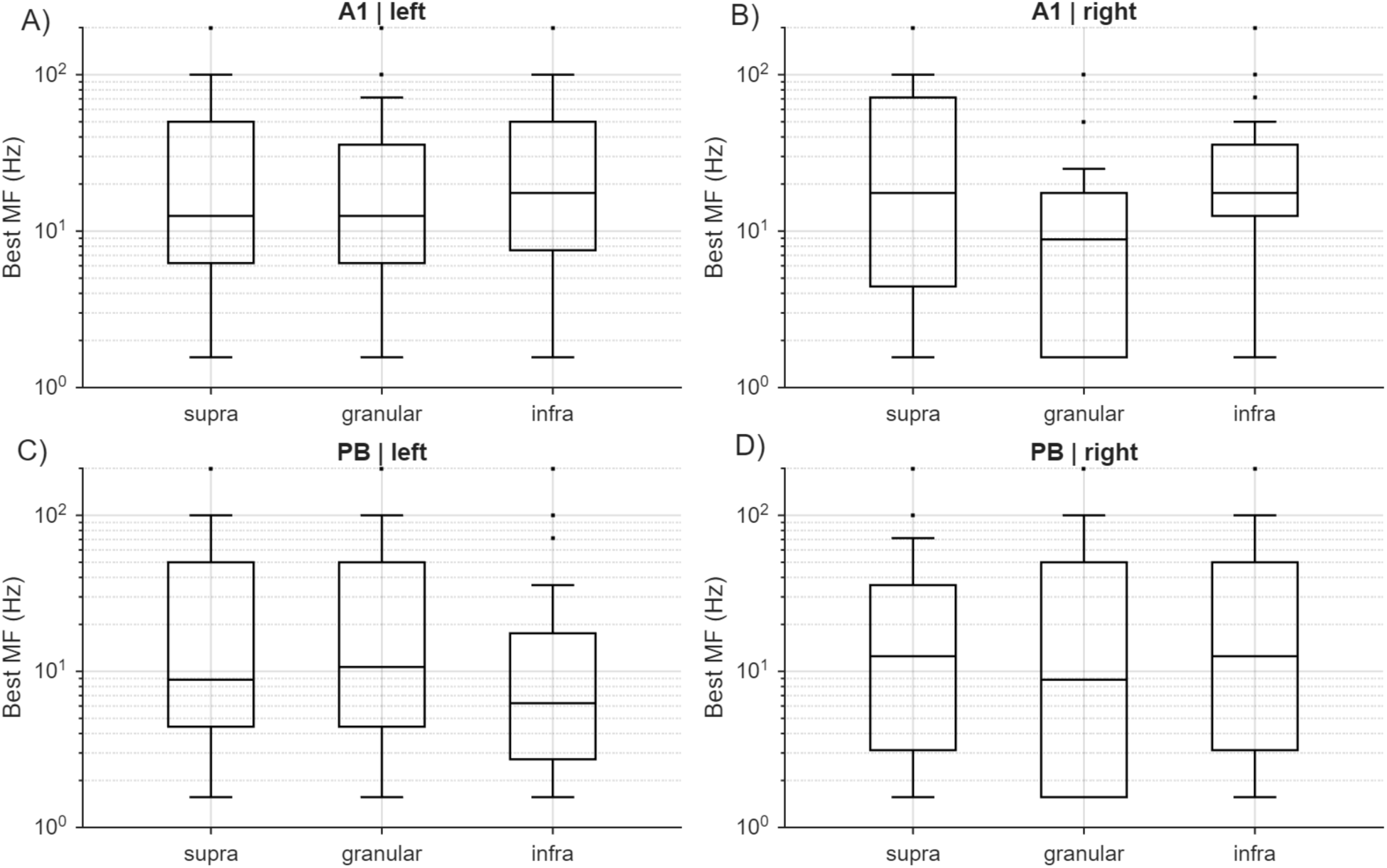
Best decoded modulation frequency in area A1 and the Parabelt. A) Boxplots display the noise modulation frequency that elicited the highest decoding accuracy for multiunit activity in the left hemisphere, Area A1, in different cortical layers (Supragranular, *n =* 171 channels; Granular, *n =* 86 channels; Infragranular, *n =* 156 channels). B) Best decoded modulation frequency in the right hemisphere, Area A1 (Supragranular, *n =* 134 channels; Granular, *n =* 47 channels; Infragranular, *n =* 74 channels). C) Best decoded modulation frequency in left hemisphere Parabelt (Supragranular, *n =* 114; Granular, *n =* 46; Infragranular, *n =* 81). D) Best decoded modulation frequency in the right hemisphere Parabelt (Supragranular, *n =* 268; Granular, *n =* 124; Infragranular, *n =* 240).

**Table 4.** Mixed effects model of best decoded modulation frequency. *Degrees of freedom =* 1534, *n = 1531* channels.

| Model: Best MF ~ Area + Layer + Hemisphere + Area*Layer + Layer*Hemisphere + Hemisphere*Area + (1\|Channel), Degrees of freedom = 1532. |  |  |
| --- | --- | --- |
| Term: | t-statistic | P-value |
| Right hemi (vs. L) | -1.5 | 0.12 |
| Parabelt (vs. A1) | 0.15 | 0.87 |
| Supra (vs. Gran) | 1.8 | 0.06 |
| Infra (vs. Gran) | 2.2 | 0.02 |
| Infra*Right Hemi | -0.41 | 0.68 |
| Infra*Parabelt | -3.5 | 0.0004 |

### AM encoding over time

Having characterized how the encoding of AM differs across regions using the entire response waveform, it was then of interest to investigate how encoding unfolds across time, which can address questions, albeit indirectly, about spike-timing vs. spike rate codes in previous studies (e.g. Hoglen et al., 2018; Yin et al., 2011; see Introduction). Analyzing decoding results over time also addresses what time-points in the stimulus response may be driving the decoding accuracy in our previous results, which utilized the entire response waveform (e.g. Figures 1-3). To address these issues, responses were binned into 10 or 50 ms bins and the decoding process was repeated for each bin (see Methods). Figure 6 shows example decoding results over time, for the 6.3 Hz AM noise stimulus for A1 (Figure 6A-B) and parabelt (Figure 6C-D). These results complement the mutual information analyses, by showing that at a low modulation frequency (below ∼25 Hz), sites in the parabelt often displayed far above-chance decoding accuracy. They also suggest that the highest magnitude response, in CSD or MUA, often encountered at the onset of the stimulus (Figure 1), does not always correspond to the peak decoding accuracy, which could be found throughout the duration of the stimulus response; i.e. maximum decoding occurred sparsely over time. Consistent with this, decoding accuracy of different modulation frequencies exhibited peaks throughout the response, from onset latency through 700 ms (Figure 6E-H) for both the CSD and MUA signals. These results also suggest that the time-varying aspect of the response to AM contributes heavily to its discriminability in both A1 and the parabelt (see Discussion). This observation likely depends on our use of population-level activity, which has been found to exhibit robust temporal encoding in a previous single-unit study (see Introduction and Discussion; Downer et al. 2021).

**Figure 6.**
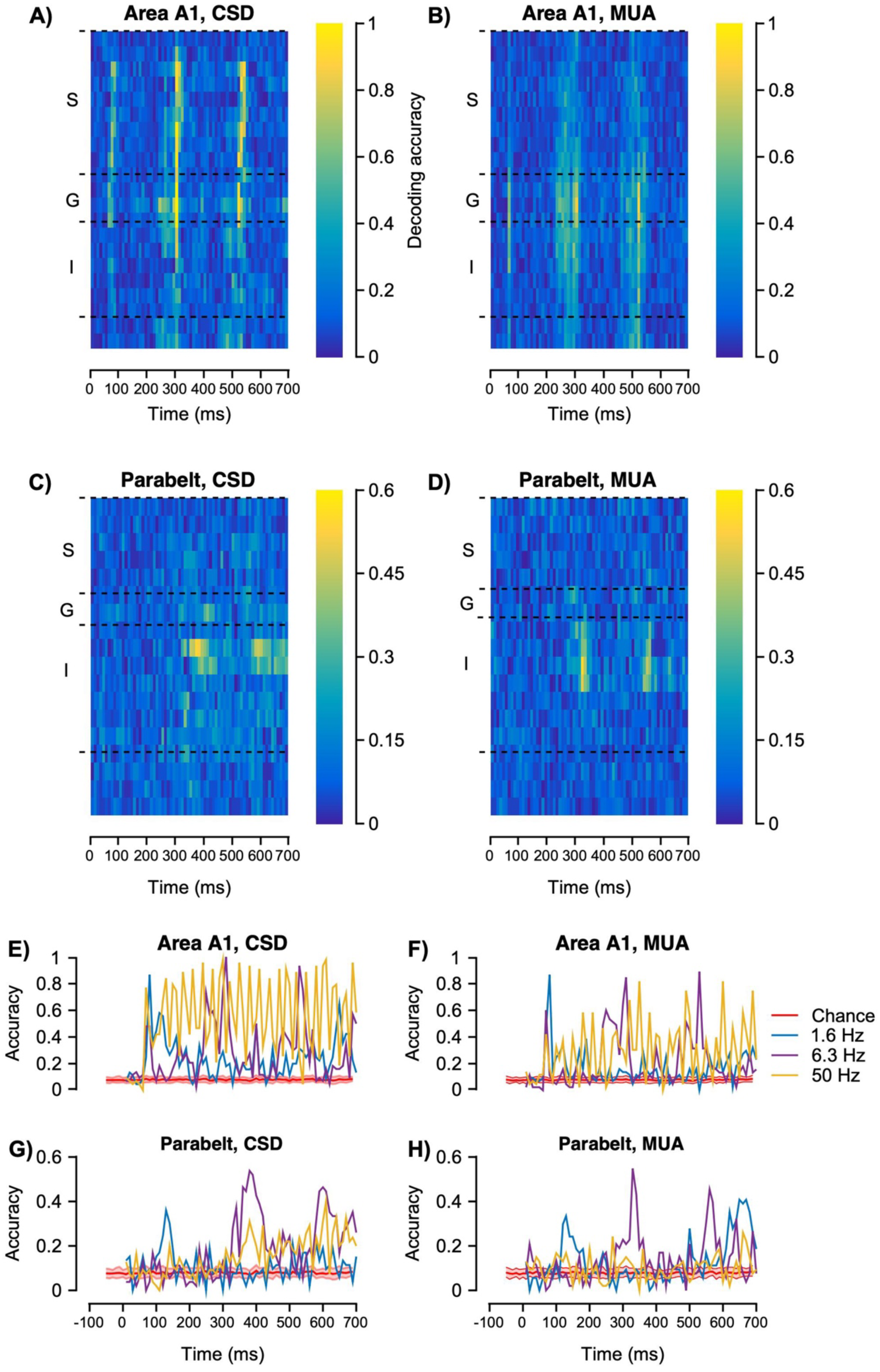
Decoding AM noise over time. **A**) Representative example of decoding the CSD signal over time, for the 6.3 Hz AM noise condition in left hemisphere area A1 (*n =* 50 trials). The decoding results for the same site’s MUA are shown in (**B**). **C**) Representative example of decoding the CSD signal over time, for the 6.3 Hz AM noise condition in the right hemisphere parabelt auditory cortex (*n =* 50 trials). The decoding results for the same site’s MUA are shown in (**D**). In all heat maps, the bin size was 10 ms. **E**) Decoding accuracy using the CSD signal from a granular layer channel (11) in panel (A) across a subset of modulation frequencies used in the study. **F**) Decoding accuracy using the MUA signal from A1 granular layer channel (11) in panel B across a subset of modulation frequencies used in the study. **G**) Decoding accuracy using the CSD signal from an infragranular layer channel (9) in panel (**C**) across a subset of modulation frequencies used in the study (1.6-200 Hz). **H**) Decoding accuracy using the MUA signal from an infragranular layer channel (11) in panel B across a subset of modulation frequencies used in the study. Chance level decoding was calculated after shuffling the decoding labels and is indicated by the red trace. Error bands represent 95% confidence intervals (1,000 permutations).

**Figure 7.**
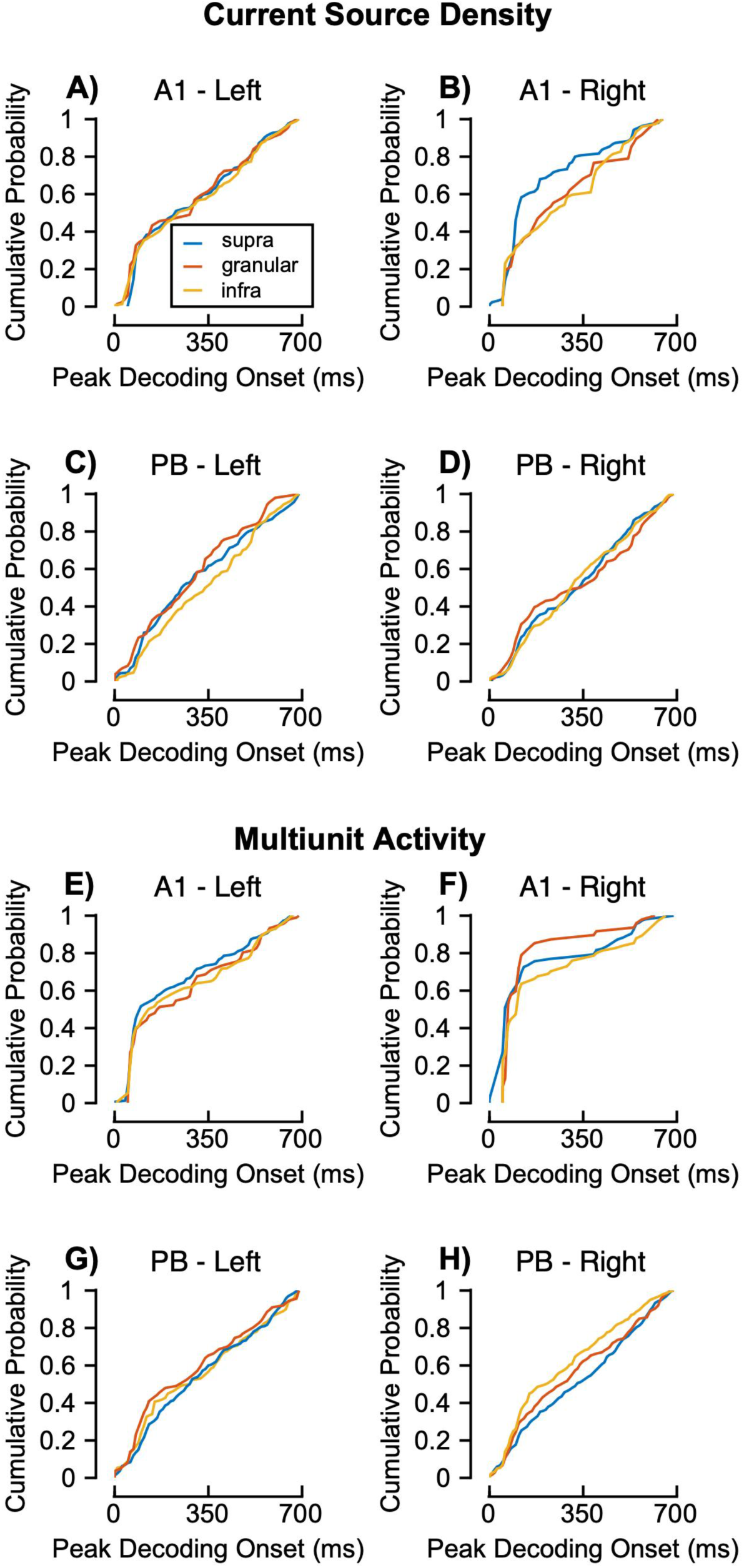
Cumulative distributions of peak decoding onset in A1 and parabelt in response to amplitude modulated noise using a 10 ms bin size. Distributions of peak decoding accuracy values, derived from the CSD signal, are depicted for left and right hemispheres of area A1 (**A-B**), and the parabelt (**C-D**). Distributions of peak values derived from the MUA signal are depicted for left and right hemispheres of area A1 (**E-F**), and the parabelt (**G-H**). Only above-chance values are displayed (see Methods).

To quantify these observations, the peak decoding time, across all modulation frequencies, was compared across layer, area, and hemisphere. The right hemisphere had significantly earlier peak times (Table 5), though this appeared to be restricted to area A1, indicated by the significant interaction of parabelt with the right hemisphere (Figure 5; Table 5). The parabelt had significantly later peak times in both the CSD and MUA signals. In the infragranular layers of the parabelt, MUA peak decoding times were earlier than predicted by the infragranular layer term on its own (Table 5). The same mixed effects model analysis revealed similar size and direction of effects for the 50 ms binned decoding results (Tables 6).

**Table 5.** Mixed effects model of peak decoding onset time in the MUA signal, using a 10 ms bin. *Degrees of freedom =* 1716 , *n = 1713* channels.

| Model: Peak Time ~ Area + Layer + Hemisphere + Area*Layer + Layer*Hemisphere + Hemisphere*Area + (1\|Channel) |  |  |
| --- | --- | --- |
| Term: | t-statistic | P-value |
| Right hemi (vs. L) | -2.6 | 0.008 |
| Parabelt (vs. A1) | 2.48 | 0.013 |
| Infragranular (vs. Gran) | 0.99 | 0.3 |
| Infragranular layer*Parabelt | -1.11 | 0.26 |
| Right Hemi*Parabelt | 2.9 | 0.004 |

**Table 6.** Mixed effects model of peak decoding onset time in the MUA signal, using a 50 ms bin. *Degrees of freedom =* 1716, *n = 1713* channels.

| Model: Peak Time ~ Area + Layer + Hemisphere + Area*Layer + Layer*Hemisphere + Hemisphere*Area + (1\|Channel) |  |  |
| --- | --- | --- |
| Term: | t-statistic | P-value |
| Parabelt (vs. A1) | -0.08 | 0.5 |
| Supra (vs. Gran) | -2.1 | 0.03 |
| Infragranular<br>(vs. Gran) | -0.1 | 0.91 |
| Supra<br>layer*Parabelt | 3.2 | 0.002 |
| Infragranular<br>layer*Parabelt | 0.8 | 0.39 |
| Right Hemi (vs.<br>L) | -1.3 | 0.2 |

## Discussion

### Parallel processing across auditory cortex

The present report builds on previous work on how sounds are encoded across the auditory cortical hierarchy, which exhibits serial (A1-Belt-Parabelt) and parallel organization (the medial geniculate body, MGB, projects to multiple subdivisions in parallel). Rather than a uniform degradation of temporal precision, the laminar gradient in encoding observed in A1 (Granular > Infragranular > Supragranular) was inverted in the parabelt (Supragranular > Infragranular > Granular; see Table 3). This difference is noted with some caution, given the general lack of significant AM encoding in the PB. The laminar gradient inversion may result from differences in nonlemniscal thalamic projections across auditory cortex. Specifically, extragranular A1, Belt, and Parabelt receive projections from the magnocellular MGB, and the dorsal MGB projects to layers 3b and 5 of Belt and Parabelt; avoiding A1 (Burton and Jones, 1976; Hackett, 2011; Huang and Winer, 2000). In the parabelt, feedforward cortical input to layer 4 from belt (Hackett, 2011; Hackett et al., 2014), in combination with parallel dorsal MGB input to L3b/5 could, theoretically, lead to the enhanced extragranular encoding we observe in the PB, relative to granular PB (Table 1). This idea is consistent with findings of rapid onset latencies, rivaling A1, observed in caudal belt, and robust temporal encoding in anterior dorsal MGB (Bartlett and Wang, 2011; Kajikawa et al., 2011). Unlike dorsal MGB, magnocellular MGB does not avoid A1 (Burton and Jones, 1976; Hackett, 2011; Huang and Winer, 2000). However, its dense projections to layer 1 are thought to position it as a modulatory influence (Jones, 2001; Lakatos et al., 2020), and our data do not indicate that it is integrated with layer 4 thalamic input to provide enhanced extragranular encoding of AM.

### Serial processing across auditory cortex

Previous NHP studies have described hierarchical progressions in stimulus selectivity that coincide with the hierarchical increase in response latency in primary, secondary, and tertiary (parabelt) auditory cortex (Camalier et al., 2012). For instance, NHP studies of AM encoding have discovered notable differences in encoding between primary and secondary auditory cortex. Namely, while primary AC encodes AM depth with an increasing rate code, secondary belt auditory cortex exhibits an opponent (increasing/decreasing) code for AM (Downer et al., 2017; Johnson et al., 2024). Consistent with this, fMRI studies have noted some degree of selectivity for longer temporal intervals in nonprimary auditory cortices, particularly in the parabelt (Dheerendra et al., 2021). An extended temporal integration window, and broad frequency tuning likely contributes to the parabelt’s enhanced response, relative to area A1, to harmonic sounds with varying degrees of temporal coherence across frequency channels, termed “auditory figure ground” stimuli (Schneider et al., 2018). Across studies of the cortical hierarchy, there appears to be a notable reduction in response to simple stimuli, such as tones, and an increase in selectivity for complex stimuli with characteristics, e.g. low modulation frequencies. This likely underpins enhanced selectivity for species-specific communication sounds, and associated visual stimuli, as one ascends the hierarchy (Mackey et al., 2024b).

### Previous laminar studies

The results presented here provide the first study of AM encoding utilizing laminar resolution at multiple levels across the NHP auditory cortical hierarchy. Previous laminar studies in cat primary auditory cortex found that the extragranular layers preferentially encode slower AM relative to the thalamorecipient granular layer (Atencio and Schreiner, 2010), and that spectrotemporal feature selectivity is hierarchical i.e. Granular > Supragranular > Infragranular (Atencio et al., 2009). This is consistent with what we observed for granular MUA in A1, though interestingly this trend was much less apparent in the parabelt (Figure 3 E-H); and Infragranular encoding was on par with Granular layer encoding in both A1 and Parabelt (Figure 3E-H). This may indicate a role of top-down infragranular outputs in the awake state. Murine studies have suggested robust differences in spectrotemporal tuning between layer 5 and layer 6 neurons, with L5 being sluggish relative to L6, and exhibiting much broader, less sparse tuning to spectral and temporal sound features (Williamson and Polley, 2019). Our measures of local population activity (CSD and MUA) are likely integrating across these populations, so, perhaps unsurprisingly, we did not see a bimodal distribution (reflecting L5 and L6 units) in infragranular AM encoding measures. However, the robust AM encoding in infragranular populations (e.g. Infragranular encoding being on par with Granular layer encoding in Figure 3E-F) is consistent with these previous data on L6 units. These studies, in combination with the present report, describe some of the only data on the intracortical circuits underlying temporal sound envelope encoding in auditory cortex. The present report now builds on this knowledge by quantifying laminar-resolved AM encoding in the NHP parabelt.

### Task engagement

Though the NHPs in the present report were not engaged in a task, previous studies have suggested that stimulus complexity and task relevance exert substantial influence on stimulus responses, particularly in nonprimary cortical areas. In secondary and tertiary auditory cortices of ferrets (the dorsal posterior ectosylvian gyrus, and rostral ventral posterior auditory field, respectively), hierarchical differences, such as the progressive influence of task-related factors, have been well-studied (Atiani et al., 2014; Elgueda et al., 2019). A similar hierarchical progression is present between primary and secondary auditory cortex of NHPs discriminating AM noise, namely, single-unit discriminability via rate and temporal codes are enhanced by task-engagement (Niwa et al., 2015). Entrainment to rhythmic stimuli also appears to be behaviorally-gated in this same sense (O’Connell et al., 2015). Thus, if our NHPs had been engaged in a task, particularly one that indexed AM feature discrimination, we would expect enhanced encoding relative to what we report here.

### Decoding studies

Previous decoding studies offer insight into the present results (Downer et al., 2021; Tovar et al., 2020). Tovar et al. (2020) leveraged multivariate pattern analysis across channels to reveal that visual gradient orientation information is relatively well preserved across all layer compartments in V1, and eye-of-origin information decreases in the infragranular layers. Tovar et al. also noted that repetition suppression, which is greatest in the extragranular layers, contributed to their enhanced decoding of stimulus repetition. This may be analogized to our results, where the granular layer often exhibited the greatest mutual information about different modulation frequencies, likely due to its rapid response characteristics. Though this pattern was most notable in area A1, and the parabelt displayed a less clear granular > supragranular > infragranular gradient in AM encoding. Another study that utilized decoding analysis that is relevant to the present results is Downer et al. (2021). Downer et al. utilized decoding analyses on simulated and empirical neuronal populations from single-unit recordings, and revealed robust population-level synchronization to a wide range of modulation frequencies, 4-512 Hz (Downer et al., 2021). This is consistent with our MUA measures in area A1. Both studies suggest code-agnostic measures (e.g. linear discriminant analysis in the present report) on local population responses can reveal information not readily apparent in single-unit measures using phase-locking or an average rate code. Such code-agnostic measures also avoid commonly noted problems with vector strength and the Rayleigh test for synchrony, namely potential for false discoveries (Yin et al., 2011), and parametric statistical assumptions (Kajikawa and Hackett, 2005).

### Interhemispheric differences

Our analysis demonstrates similar hierarchical and inter-hemispheric differences to human subjects in Boemio et al. (2005), and our results resemble previous NHP studies demonstrating sound-driven hemispheric asymmetries (left hemisphere enhancement) in the lateral sulcus, and the extent of the superior temporal gyrus, including the temporal pole (Poremba et al., 2004, 2003). Our results suggest the supragranular layers are the main contributing layer to the left-hemisphere dominance in encoding a wide range of temporal fluctuations (1.6-200 Hz), which aligns with other studies which defined supragranular sources potentially underlying noninvasive measures in fMRI, EEG, and MEG studies (e.g. (Kajikawa et al., 2024; Lakatos et al., 2020; Leszczyński et al., 2020; Peterson et al., 1995)). Additionally, a previous NHP study has similarly demonstrated enhanced entrainment of low frequency (∼1.6 Hz) oscillations in left, relative to right, supragranular A1, which corroborates our results (O’Connell et al., 2015).

## Acknowledgements

Funding sources: R01DC015780, R01DC012947 and R01DC019979.

The authors thank Dr. Monica N. O’Connell for helpful comments on an earlier version of the manuscript.

## Notes

### Competing Interest Statement

The authors have declared no competing interest.

### Summary of Updates

Revised statistics, described in the abstract, results, and discussion.

